# Antiviral Resistance against Viral Mutation: Praxis and Policy for SARS-CoV-2

**DOI:** 10.1101/2021.02.25.432861

**Authors:** Robert Penner

## Abstract

New tools developed by Moderna, BioNTech/Pfizer, and Oxford/Astrazeneca, among others, provide universal solutions to previously problematic aspects of drug or vaccine delivery, uptake and toxicity, portending new tools across the medical sciences. A novel method is presented based on estimating protein backbone free energy via geometry to predict effective antiviral targets, antigens and vaccine cargos that are resistant to viral mutation. This method, partly described in earlier work of the author, is reviewed and reformulated here in light of the recent proliferation of structural data on the SARS-CoV-2 spike glycoprotein and its latest mutations in the variants of concern and several further variants of interest including all international lineages. Particular attention to structures computed with Cryo Electron Microscopy allows the novel approach of probing the pH dependence of free energy in order to infer function. Key findings include: collections of recurring mutagenic residues occur across strains, presumably through cooperative convergent evolution; the preponderance of mutagenic residues do not participate in backbone hydrogen bonds; metastability of the spike glycoprotein limits the change of free energy from before to after mutation and thereby constrains selective pressure; and there are mRNA or virus-vector cargos which target low free energy peptides proximal to conserved high free energy peptides providing specific recipes for vaccines with greater specificity than the current full-spike approach. These results serve to limit peptides in the spike glycoprotein with high mutagenic potential and thereby provide *a priori* constraints on viral and attendant vaccine evolution. Scientific and regulatory challenges to nucleic acid therapeutic and vaccine development and deployment are finally discussed.

## 1. Introduction

Breakthrough capabilities for cellular delivery of engineered nucleic acids have overcome major hurdles [1]. A critical question is which nucleic acid cargos to deliver for what effect. Other aspects, such as cellspecific uptake or translation promoters, will surely be further refined. One can anticipate decades of immunological and protein placement and replacement therapeutic advances [2].

First applications have been the deployment of SARS-CoV-2 vaccines, whose cargo is sensibly given by nucleic acid from the virus itself. Both the Moderna and BioNTech vaccines deliver mRNA for the full spike glycoprotein, albeit a prefusion stabilized mutation K986P/V987P, called 2P, patented in 2016 in the general context of *β*-coronaviruses, while the adenovirus-vectored vaccines developed by Oxford and others deliver DNA instructions for the full wild-type spike.

We have been fortunate so far in several regards: the 2P mutation was already known; effective reverse translation from protein to nucleic acid cargo was already solved; the derived antibodies are broadly neutralizing, including penetrating glycan shielding; and the inevitable viral mutations are only recently partly escaping vaccine efficacy [3].

Several Variants of Concern (VoCs) have arisen from the classical Wuhan strain, notably the D614G mutation, which achieved global prevalence, the U.K. strain B.1.1.7, the South African strain B.1.351 and the Brazilian strain P.1. Several other Variants of Interest (VoIs), namely, the U.K./Rwanda strain A.23.1, the S. Calif. strain B.1.429, the U.K./Denmark/U.S.A./Nigeria strain B.1.525, the New York strain B.1.526, and the emerging Maharashtra strains B.1.617 and B.1.618 are studied here. The mutations characterizing these various strains are annotated in [4] and explicated in SI Table 2, along with those of the so-called Internatinal Lineages B.1.1.7, B.1.351, P.1, A.23.1, B.1.429 and B.1.525. The characterizing mutations of the VoCs are illustrated in Fig. 1.

**Figure 1.**
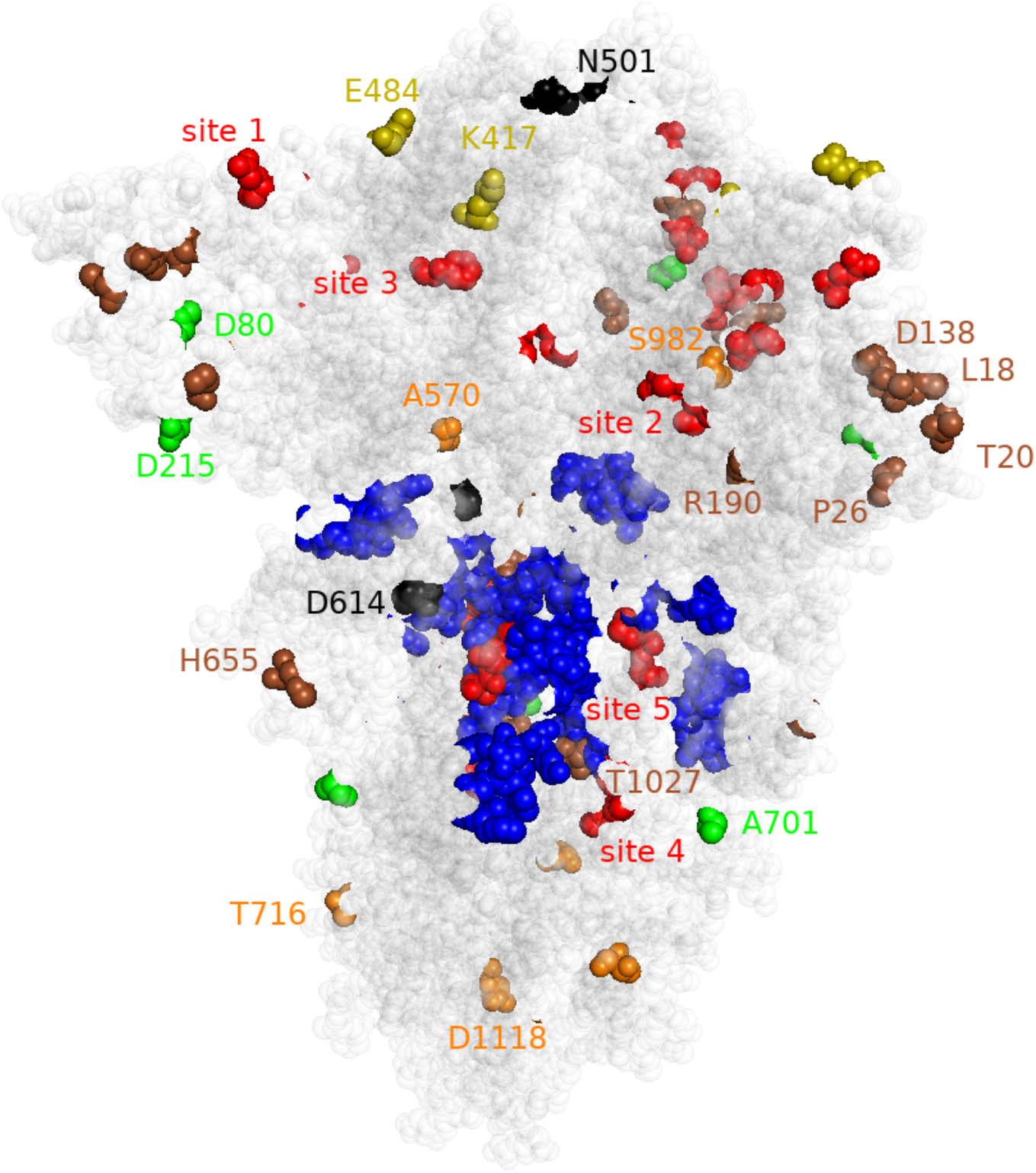
Mutated sites of the SARS-CoV-2 spike glycoprotein for the variants of concern depicted in PDB file 7df3, which does not model residue P681. Orange, brown and green residues are respective mutations for the U.K., Brazil and S. Africa variants, with olive residues K417N and E484K common to the latter two, and black residues N501Y and D614G common to all three. Also depicted are red/blue residues for the active/passive sites of interest explained in the text.

Recall (cf. [5]) that the intracellular state is communicated to the extracellular adaptive immune system through several pathways including the major histocompatibility complexes (MHCs) and through intact-antigen presenting dendritic cells. These complex processes are too much to review here, culminating in the eventual production of antibodies and memory cells through B- and T-cell clonal expansion, with both fortuitously provoked by nucleic acid vaccines.

Several points deserve amplification. Full-molecule cargos, as have been implemented thus far, cannot be certain to target neutralizing epitopes, especially for a pathogen with high morbidity. Though other properties pertain, it is the protein geometry that is the driving force for immunological recognition and response, in intact-antigen display and in the proteolytically derived peptides for MHC presentation, so any vaccine cargo in isolation must determine a protein that has a similar three-dimensional folded structure to that in the full molecule. Viral evasion can include decoy non-neutralizing epitopes. Viral mutation selecting for favorable traits is especially brisk for RNA viruses such as SARS-CoV-2, in principle thwarting antiviral strategies. One key finding here is that there are *a priori* constraints upon mutagenicity of a residue, such as participating in a backbone hydrogen bond (cf. next section), suggesting that multivalent mRNA vaccines could be developed which are universal for families of likely viral variants.

## 2. Materials and Methods

See [6, 7, 8] for further details on this overview of materials and methods. The atoms in a protein peptide group lie in a plane containing the unit displacement vector of the peptide bond. A backbone hydrogen bond (BHB) from N-H to O=C, where carbon C and nitrogen N lie in the protein backbone, therefore determines a pair of such planes containing vectors, ordered from hydrogen bond donor to acceptor. There is a unique rotation of space carrying the first plane and the displacement vector within it to the second and preserving the natural orientations on these planes. It follows that a BHB uniquely determines a rotation of space.

A suitably unbiased and high quality subset called HQ60 (for “high quality at 60% homology identity”) of the Protein Data Bank (PDB) therefore determines the histogram of all such rotations for all the constituent BHBs as in [8]. This distribution on the space of rotations, i.e., this histogram 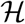 (depicted in SI Fig. 1) representing the 1.2 million or so rotations of BHBs of the proteins in HQ60, regarded as representative of all proteins, provides a fundamental database 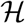, which allows the approximation of free energy, as follows.

A standard tool in protein theory is the so-called Pohl-Finkelstein quasi Boltzmann Ansatz [9, 10, 11], which allows the estimation of BHB free energy (BFE) of a residue in any protein based upon this database, where in effect low density in the histogram 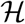 corresponds to high free energy. By definition, the BFE of a residue is the maximum of the free energy of all BHBs between the C=O or N-H nearest it along the backbone. This method of estimating backbone free energy is a new tool employed here that should be of general utility across structural biology.

Specifically, one decomposes the space of rotations into small uniform cubes, and takes the density of 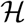 in each cube to define a real-valued piecewise-constant function *h* on the space of rotations. The entire distribution 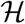 is strongly unimodal, with the mode *μ* (i.e., the point of highest density in 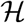) occurring at the rotation corresponding to an internal turn of an ideal (left) *α* helix. Now given any new subject rotation *ρ*, such as that corresponding to a BHB in the spike glycoprotein of our central interest, define

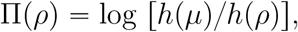

namely, the negative of the logarithm of the difference of the probabilities of the subject rotation compared to the ideal *α* helix, according to the quasi Boltzmann Ansatz. The units are actually given in *kT_C_*, where *k* is the Boltzmann constant and *T_C_* is an effective temperature, the so-called conformational temperature (which is approximately the protein melting temperature), cf. [10]. As an approximation for absolute BFE by taking actual temperature for conformational temperature, we have simply Π(*ρ*) — 2.9 in units of kcal/mole, as argued in [6], where the rotation *μ* for the ideal *α* helix has BFE of −2.0 kcal/mole.

Viral glycoproteins, for which there is known to be large conformational activity during both receptor binding and fusion, are studied as a case in point. The basic thesis proven in [6] is that if the BFE of a residue lies in the 90th percentile, i.e., exceeds 4.6 kcal/mole, then within one residue of it along the backbone during molecular activation, the sum of the two adjacent backbone conformational angles changes by at least 180 degrees. (This sentence corrects an unfortunate earlier misstatement of the precise result.) In short: large BFE implies large backbone conformational change. The converse is not valid.

The methods of [6] are applied here to the structure files from [12–[20] annotated in SI Table 1. These high quality structures represent uncleaved and antibody- and receptor-free SARS-CoV-2 spikes computed by Cryo Electron Microscopy, which allows for control of the pH of the sample through specified pH of the buffer at a variety of values, ranging from the acidic 4.5-5.5 of the endocytic pathway, to the 7.0-7.5 of the extracellular environment, and to the extremely basic 8.0, occurring for mammals only in the mitochondrial matrix. Several structures include various point mutations including 2P for prefusion stabilization, as noted in SI Table 1, as well as the globally prevalent D614G in the last 10 structures in the list. Note that for such stabilized glycoproteins, the approximations here give only a rough idea of BFE for the native molecule *in vivo* at low pH.

## 3. Results

Certain considerations raised in the Introduction can be addressed through protein backbone free energy. The methods of [6] are applied to *α*- and *β*-coronavirus spike glycoproteins in [7], leading to five conserved so-called active sites of interest, depicted in red in Fig. 1 and described by (donor, acceptor, acceptor) triples of residues comprising high BFE bifurcated BHBs, namely, (C131, L117, Q134), (I203, V227, D228), (F392, V524, C525), (M1029, V1034, L1035) and (G1059, S730, M731), which are conserved across seven human coronavirus spikes aligned using both high BFE and amino acid sequence, a presumptive proxy for functional alignment. The idea is that high BFE is evolutionarily conserved only in case of functional dependence, and conservation across different coronaviruses implies critical function conserved also across different strains of the same coronavirus, i.e., they provide vaccine or therapeutic targets that should be universal across strains.

Since the computations of [7] were performed before there was ample data on the SARS-CoV-2 spike in the PDB, the first consideration here is confirmation that the active sites of interest remain so for the more recently considered structures from SI Table 1. This is indeed the case with the following stipulations: low pH <6.0 disrupts bifurcated hydrogen bond high BFE especially for sites 1,2 and 3; linoleic acid binding at pH 7.0 disrupts this bifurcated high BFE of all five sites; for site 1 even at high pH ⩾ 7, the mutation R685S disrupts bifurcated high BFE, and D614G shifts it to nearby residues; the structure file 6×29 at pH 7.4 inexplicably has all five bifucations disrupted.

These five conserved and critical active sites of interest are presumably promising antiviral targets but are unsuitable as antigens since high BFE peptides cannot fold in isolation as they do in the full molecule, requiring low free energy scaffolding for metastability. This leads to consideration of low BFE sites as targets and unlikely mutational loci. Low BFE peptides likely fold in isolation as they do in the full molecule and thus provide effective targets. Moreover by definition [10, 6], low BFE geometry is stabilized by large numbers of peptides, so mutations should preserve the geometric structure of the antigen; most mutations that occur in these regions are thus not critical and will not be selected by evolutionary pressures, though on the other hand, charge or other residue attributes might lead to positive selective pressure. The question thus becomes whether there are low BFE sites, so-called passive sites of interest, whose immobilization might interfere with these five active sites nearby.

In fact, there are four passive sites of interest common to all structures, as enumerated in Table 1 and illustrated in blue in Fig. 1. All sites occur on the molecular surface, and sites 2, 3 and 4 are nearby one another in space and combine into one supersite. Coding mRNAs for these sites should provide salutary nucleic acid vaccine cargos, as has already been discussed. These passive sites of interest all occur in *α* helices, sometimes with other secondary structure motifs at their ends. Note that *α* helices are not necessarily low BFE, and low BFE occurs elsewhere beyond *α* helices.

**Table 1.** Average BFE in kcal/mole and percentage of residues with negative BFE for each site computed without taking into consideration hydrogen bond donors in the first three or acceptors in the last three residues of each site.

|  | <u>site 1</u> | <u>site 2</u> | <u>site 3</u> | <u>site 4</u> |
| --- | --- | --- | --- | --- |
| Residues | 295-304 | 816-825 | 949-962 | 1001-1028 |
| Average | 0.00 | 0.14 | -1.16 | -1.60 |
| Percent | 51.83 | 55.73 | 82.61 | 86.64 |

Many among the mutant residues are *missing* from most of the structures in SI Table 1, cf. SI Table 2. One might infer that these residues lie in disorganized regions of the protein, or perhaps that the experiments were less well controlled there. As an indication that the former pertains, one computes that in the specified collection of structures with an average resolution of 3.02 Angstrom, on average, the clash score is 2.1, Ramachandran outliers amount to 0.21 percent and sidechain outliers amount to 0.79 percent, where the last datum is relatively insignificant here since the BFE depends only upon the protein backbone and its adjacent C=O.

Further analysis (not presented), taking large/small B-factor to imply disorganization/order of these structures, shows that VoC mutated residues which are present in structures are highly disorganized in S1 and ordered in S2. Moreover, the active sites of interest in S1 are not highly disorganized and in S2 are highly ordered, while the passive sites of interest, all of which lie in S2, are also highly ordered. This is in keeping with the general trend in the prefusion spike that the membrane-distal molecular surface of S1 is largely disorganized and S2 is ordered.

Given a residue number *n*, let *D* = *D_n_* (for *disorganized*) denote the number of PDB files, among the 21 native structure files (not allowing D614G, so omitting the last 10 files from SI Table 1) from which *n* is missing, and let *U* = *U_n_* (for *unbonded*) be the number of PDB files among the 21 where residue number *n* is not missing, but also does not participate in a BHB, in the sense that the C=O and N-H nearest to the C^*α*^ of the residue along the backbone do not participate in a BHB. Thus, *D_n_* + *U_n_* is the number of PDB files in which residue number *n* in any case does not participate in a BHB.

One key analytic finding of this paper is that *D_n_* + *U_n_* is large for most of the mutagenic residues *n* among the Internatinal Lineages, cf. SI Table 2. Namely, *D_n_* + *U_n_* > 16 out of the full 25 mutagenic residues in the Internatinal Lineages in SI Table 2; the averages of *D/U* over all residues in the entire spike molecule are 1.5/5.2, respectively, and the average BFE is 2.1 kcal/mole.

Another key observation is the evidence of convergent evolution in SI Table 2, via a mix-and-match of residue mutations. Among these are: the predominant D614 has modest positive BFE presumably balanced by 614G so as to preserve overall metastability; E484K, having *D/U* = 5/5 common to five strains, with the interesting co-occurrent interplay with S477N, having *D/U* = 5/8, where S477 moreover lies adjacent to E484 in the molecular surface of the RBD; P681H or P681R, having *D/U* =13/2, occur across three strains, which suggests that the charges of the Histidine and Arginine side-chains may enhance viral function; the L452R of B.1.429 compares with the L452N of B.1.617, having *D*/*U* = 0/12, where the former strain has apparently dissipated from circulation in Southern California while the latter rages in India; one concern is that the Brazilian strain with its existing E484K in common with India has also now acquired the L452R mutation, which could explain recent similarities between these evolving disease profiles to include acute cases among younger patients; the deletion Y144Δ in B.1.1.7, having *D/U* = 14/0, compares to the deletion of Y145-H146 in B.1.618 as antibody escape techniques in the NTD; N501Y has *D*/*U* = 0/15 and occurs across multiple VoCs.

## 4. Discussion

### 4.1. Variants of Interest and International Lineages

BFE as a function of pH is plotted for the Internatinal Lineage mutant residues in Fig. 1. Residues S982 and T1027 have negative or negligible positive BFE, and R190, D1118, D614 have modest positive BFE in all cases. Residues 417 and 701 have the same peculiar character: out of the 21 PDB files of the Internatinal Lineages, 13 for residue 417 and 15 for residue 701 have at least two of the three chains of the trimer spike unbonded, and the remaining one participates in a BHB; the averages are thus not comparable to those in Fig. 2, which typically have one bond for each monomer in the trimer. Residue 888 alone is generally of high BFE, has two peaks at very high BFE for pHs 4.5 and 7.2, and a negligible positive BFE in the valley at pH 5.5, suggesting an on-off-on switch along the acidifying progression from extra- to intracellular. Residue D1118 has no bond at pH below 5 and high BFE at higher extracellular pH, suggesting possible conformational activity during binding. For up/down conformations of the receptor binding domain (cf. [7]) at pH>7.0, residue 614 has respective average BFE 1.5/3.0 for Aspartic Acid and 2.6/3.2 for Glycine, consistent with the D614G mutation affecting up/down equilibrium [20] and increasing viral transmissibility [15].

**Figure 2.**
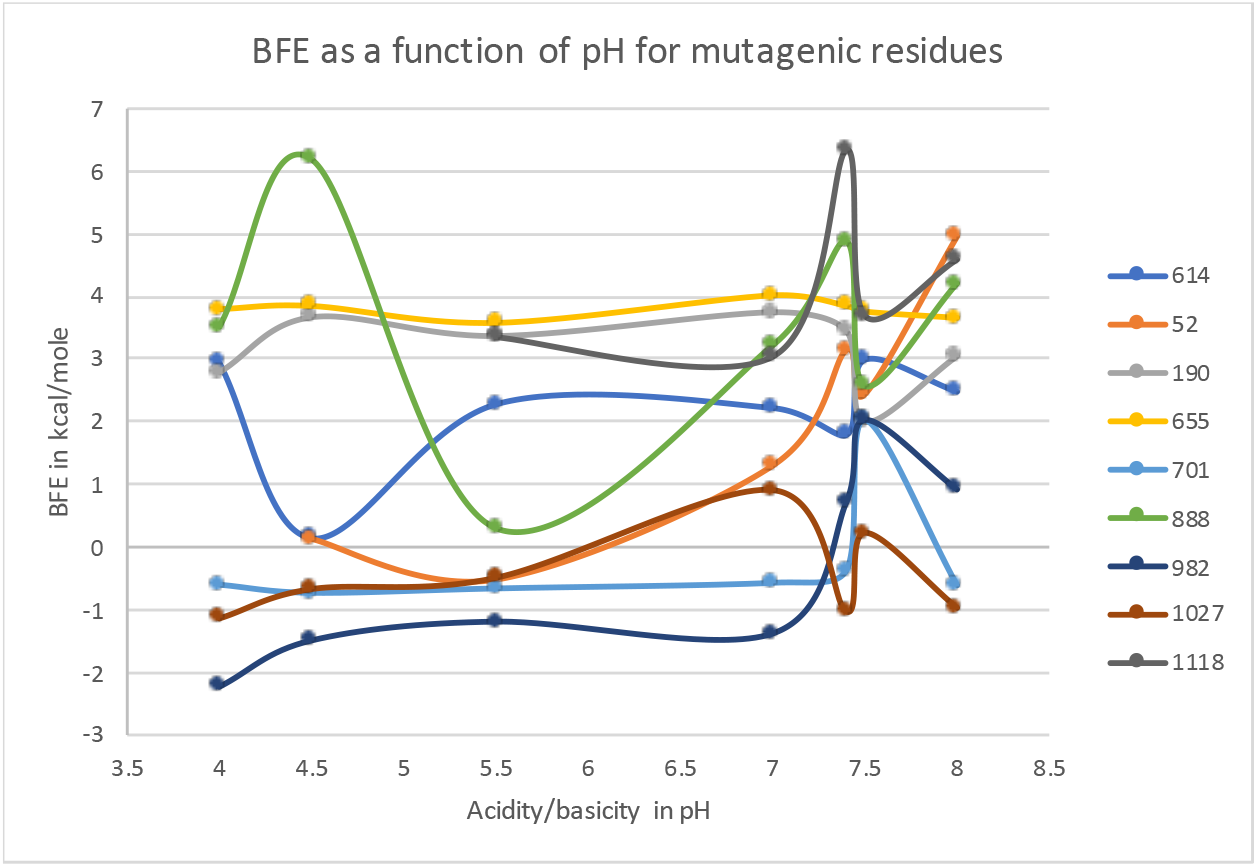
Backbone free energy as a function of pH for the bonded mutant residues in the Internatinal Lineages.

Consideration of B.1.525 (which shares the E484K mutation with B.1.351 and P.1), B.1.526, A.23.1, and B.1.429 leads to scrutiny of their additional mutated residues, annotated in SI Table 2. Residue V367, which is mutated to 367F in A.23.1, is unbonded in 15 structures with no patten for BFE in pH, and D614G leads to bonding with negative BFE, apparently of correct magnitude or in concert with other residues so as to preserve metastability.

Most interestingly for its proximity to D614, residue Q613, which is mutated to 613H in A.23.1, is bonded in all 31 structures and exhibits small positive BFE with no pattern in pH, just as for 614, but with up/down average BFE 2.0/1.7 for Aspartic Acid and with 2.8/2.2 for Glycine at 614. This opposite trend at 613 from 614 for the D614G mutation suggests similar functional changes for the two mutations with compensatory changes in BFE.

Likewise of interest, residue L452, which is mutated to 452R in B.1.429 and in the Maharashtra strain B.1.617, is present in all 31 structures of SI Table 1 with *D*/*U* = 0/12, is unbonded at pH below 7.4 (with 1 exception in file 7ad1 at pH 7.0) and at pH 7.5, and is always bonded at pH 8 with a large average BFE of 5.87 kcal/mole that is maximal in most examples. For pH 7.4, this residue is either unbonded (in 6×2c and 6×29) or again bonded with a large average BFE of 4.37 kcal/mole (in 6×2a and 6×2b).

Thus, for B.1.429, all mutated residues are disorganized or unbonded *except residue 452*, which is always present and unbonded at neutral or low pH and bonded with extremely high BFE at pH 8, which is unlikely to be coincidental. This high pH occurs in mammals only within mitochondria, which have recently been implicated in Covid-19 infection [21]. More likely though, this mutation L452R in the RBD either facilitates binding or escapes several monoclonal antibodies [22], which may be linked to the charged side-chain of Arginine.

For variant B.1.526, the mutations E484K and S477N, both of which are reported in [23] as escape mutants, are approximately equally frequent and not mutually exclusive. The BFE of residue 477 is strikingly similar to 484: missing, so presumably disorganized, in 20 structures (19 in common with 484), unbonded in 7 (5 in common with 484), and with nearly the same BFE when non-acidic. Residue T95 is unremarkable: present in all structures, unbonded in 3, and with moderate average BFE of 2.64 kcal/mole when bonded. However, the adjacent residue 96 is quite special in that it is unremarkable at pH below 8, while at pH 8, and only at this high pH, it is bifurcated with maximum BFE in at least two chains, and often all three, possibly suggesting another on-off switch.

### 4.2. Conclusions

It has already been argued that residues of high BFE are unlikely to mutate, since they would alter geometry and impede function, and of low BFE may randomly mutate but are unlikely to be selected by evolution, since geometry and hence function would be largely unchanged. Another constraint of this general nature is that mutations must approximately preserve BFE perhaps in concert with other residues: removing excessive positive or negative BFE disturbs the molecular balance required for metastability

The middle ground of no backbone hydrogen bonds and hence no BFE remains. These are precisely the residues with D+U large, which predominate in SI Table 2.

Among the remaining VoC mutant residues, three have negative free energy and three modest positive free energy, evidently with neither regime changing so substantially as to disrupt molecular metastability. Possible function can be inferred when the BFE depends upon pH.

These are general lessons that apply to any virus. For SARS-CoV-2, it has already been argued that the existing full spike molecule vaccine target may become less than optimal in light of mutations or potentially increased morbidity. Specific passive sites of interest are proposed as an alternative, and mRNA or virus-vectored vaccine could be quickly developed targeting these sites.

As a proxy for an arbitrary virus, the current approach for SARS-CoV-2 mutation is clear: keeping all other aspects fixed, substitute the spike of the variant strain for the earlier spike, whether 2P-stabilized or not. One might even sensibly deliver multi-variant cargo. In order that vaccine evolution keep pace with viral mutation, new and expedited regulatory pathways are required for the identical delivery of nucleic acid for the same protein from different variants. These remarks also apply to the targeted vaccine cargos proposed here, though the precise regulatory specifications would be more subtle.

## Supplementary Information

Figure 1, Histogram 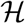 of rotations of BHBs in HQ60; Table 1, Protein Data Bank structure files upon which this work is based; Table 2, Backbone free energy of mutations characterizing various strains.

## Institutional Review Board Statement

Not applicable.

## Informed Consent Statement

Not applicable.

## Data Availability Statement

The methods of this paper are implented online from an uploaded PDB file at https://bion-server.au.dk/hbonds/

## Abbreviations

BFE: backbone free energy
BHB: backbone hydrogen bond
MHC: major histocompatibility complex
VoC: variant of concern
VoI: variant of interest.

---

**Figure 1. (Supplementary Information.).**
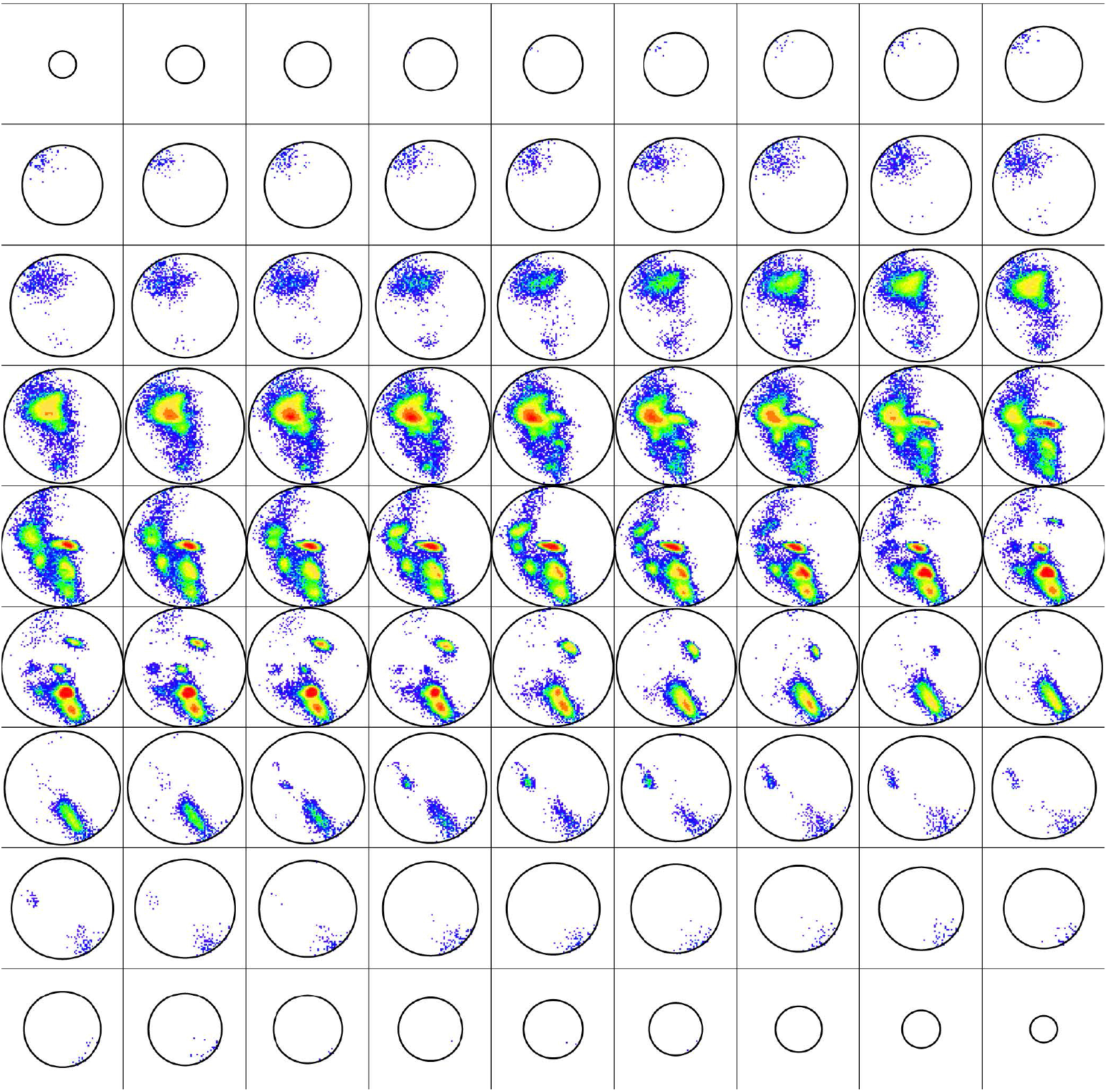
Histogram 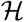 of rotations of BHBs in HQ60. The collection of all rotations of space can be represented as the ball of radius π (with each pair of antipodal points identified with one another, what mathematicians call “real projective 3-space” or “the Lie group SO(3)”), where the point in the ball has angular spherical coordinate given by the direction of the axis of the rotation, and it has radial coordinate given by the amount of rotation about the axis of rotation in radians (negative for clockwise and positive for counter-clockwise). Be that as it may, presented here are 81 horizontal slices of the histogram 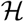 of rotations of backbone hydrogen bonds (BHBs) in HQ60 in this ball from north to south pole colored by population density from [Penner et al., *Nature Communications*, 2014], where the R-Y-G-B color is linear in the density ranging from 19,000 to 1. The mode of 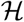 occurs at the rotation of the ideal left *α* helix in the fourth row from the top and the fourth column from the left.

**Table 1. (Supplementary Information.).**
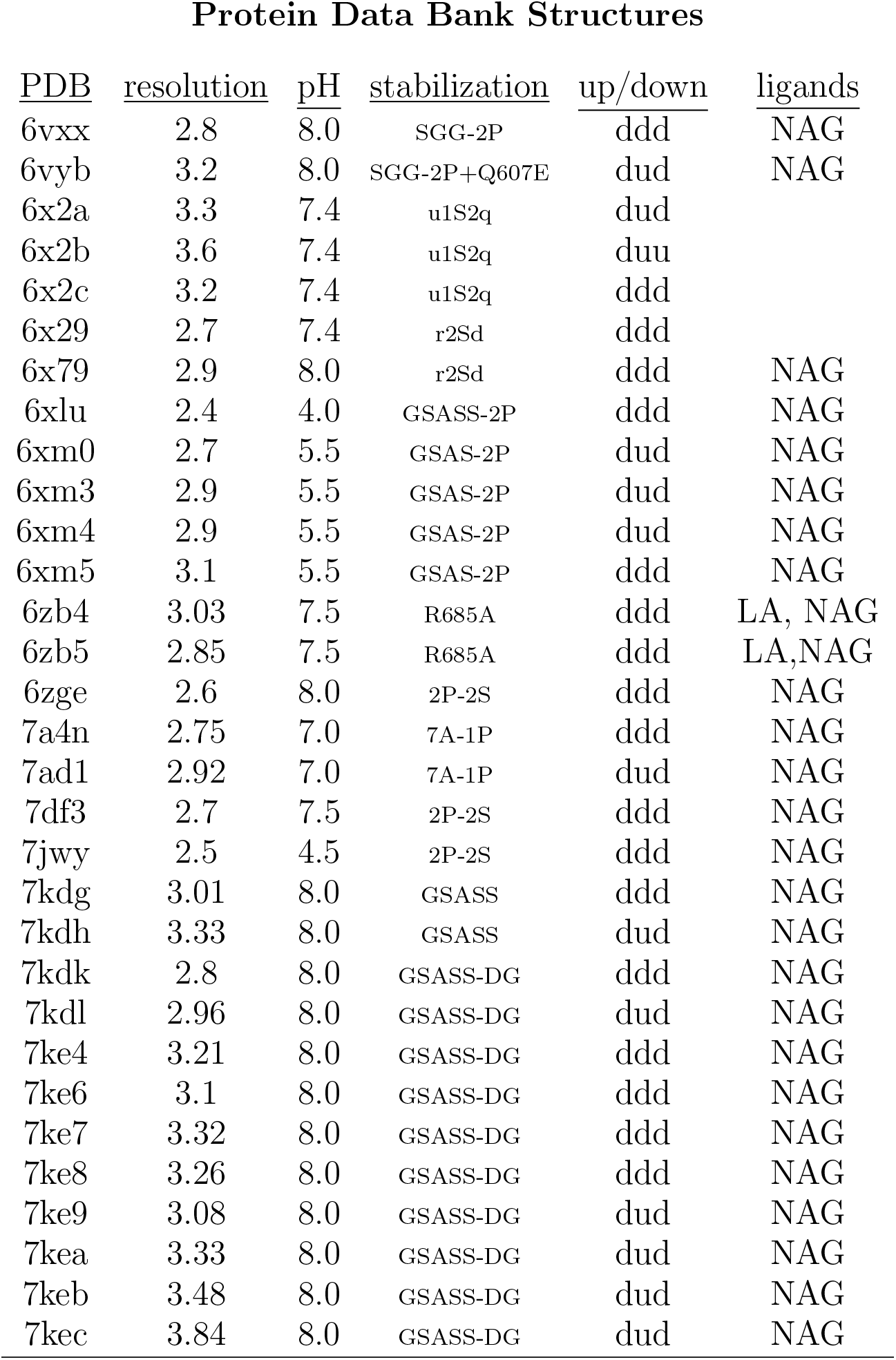
For each PDB file is given the resolution in Angstrom, the pH, the up/down conformation of the receptor binding domain for respective chains ABC, the ligands NAG= N-Acetylglucosamine or LA=linoleic acid, and the engineered mutations for stabilization, where **SGG-2P**: R682S, R683G, R685G, K986P, V987P; **ulS2q**: A570L, T572I, Q607E, R682G, R683S, R685S, F855Y, N856I, K986P, V987P; **rS2d**: S383C, R682G, R683S, R685S, D985C, K986P, V987P; **GSASS-2P**: R682G, R683S, R685S, K986P, V987P; **2P-2S**: K986P, V987P, R682S, R685S; **7A-1P**: D614N, R682S, R685G, A892P, A942P, V987P; **GSASS**: R682G, R683S, R685S, K986P, V987P; **GSASS-DG**: R682G, R683S, R685S, D614G. The final 10 files, and only these, contain the globally pervasive mutation D614G.

**Table 2. (Supplementary Information.).** Strains are Pango lineages [Rimbaut et al., *Nature Microbiology*, 2020]. The listed residues describe the mutations characterizing all current International Lineages B.1.1.7, B.1.351, P.1, A.23.1 and B.1.525, together with those residues common to those characterizing B.1.429, B.1.526 and B.1.617; the latter is the Maharashtra strain, which is of current serious concern, along with its variants that include also D618G and deletion of one or more of residues 145-146. Locations: SP=signal peptide, NTD=N-terminal domain, RBD=receptor binding domain, RBM=receptor binding motif, SD=subdomain, HR=heptad repeat, CH=central helix, CD=connecting domain; each is sometimes followed by a domain number or letter. D/U/A is either D/U, where D is the number of PDB files (out of 21 total, i.e., those in SI Table 1 which do not contain D614G) where the residue is disorganized and U the number where the residue is unbonded, or D/U/A is the BFE averaged over the various pH values provided D+U⩾11. * denotes residues out-of-range for almost all PDB files, Δ indicates deletion, and “g” indicates that Glycine is presumed to occur often, given the global prominence of D614G. B.1.526 has roughly half-half E484K and S477N, not necessarily both.

| <u>residue</u> | <u>location</u> | <u>D/U/A</u> | <u>B.1.1.7</u> | <u>B.1.351</u> | <u>P.1</u> | <u>A.23.1</u> | <u>B.1.429</u> | <u>B.1.525</u> | <u>B.1.526</u> | <u>B.1.617</u> |
| --- | --- | --- | --- | --- | --- | --- | --- | --- | --- | --- |
| L5 | SP | * |  |  |  |  |  |  | F |  |
| S13 | SP-A | * |  |  |  |  | I |  |  |  |
| L18 | SP-A | * |  |  | F |  |  |  |  |  |
| T20 | SP-A | * |  |  | N |  |  |  |  |  |
| P26 | SP-A | * |  |  | S |  |  |  |  |  |
| Q52 | NTD-A | 1.5 |  |  |  |  |  | R |  |  |
| H69 | NTD-A | 4/7 | $\Delta$ | | | | | | | |
| V70 | NTD-A | 14/0 | $\Delta$ | | | | | | | |
| D80 | NTD-A | 3/16 |  | A |  |  |  |  |  |  |
| D138 | NTD-A | 2/14 |  |  | Y |  |  |  |  |  |
| Y144 | NTD-A | 14/0 | $\Delta$ | | | | | | | |
| F157 | NTD-A | 10/4 |  |  |  | L |  |  |  |  |
| R190 | NTD-A | 3.2 |  |  | S |  |  |  |  |  |
| D215 | NTD-A | 2/17 |  | G |  |  |  |  |  |  |
| V367 | RBD-B | 0/11 |  |  |  | F |  |  |  |  |
| K417 | RBD-B | 0/13 |  | N |  | T |  |  |  |  |
| L452 | RBM-B | 0/12 |  |  |  |  | R |  |  | N |
| S477 | RBM-B | 5/8 | | | | | | | $\frac{1}{2}$ N | |
| E484 | RBM-B | 5/5 | | K | K | | K | K | $\frac{1}{2}$ K | |
| N501 | RBM-B | 0/15 | Y | Y | Y |  |  |  |  |  |
| Q613 | SD2-D | 1.7 |  |  |  | H |  |  |  |  |
| D614 | SD2-D | 2.0 | G | G | G |  | g | g | g | g |
| H655 | SD2-D | 3.8 |  |  | Y |  |  |  |  |  |
| P681 |  | 13/2 | H |  |  | R |  |  |  | H |
| A701 |  | 0/15 |  | V |  |  |  |  | V |  |
| T716 |  | 0/13 | I |  |  |  |  |  |  |  |
| F888 |  | 3.6 |  |  |  |  |  |  | L |  |
| S982 | HR1 | -0.1 |  | A |  |  |  |  |  |  |
| T1027 | CH | -0.3 |  |  | I |  |  |  |  |  |
| D1118 | CD | 4.2 | H |  |  |  |  |  |  |  |

## References

[1] P. Ball, What the lightening-fast quest for covid vaccines means for other diseases, Nature, 589, 7840, 16–18, 2021, Springer-Nature

[2] N. Pardi et al., mRNA vaccines-a new era in vaccinology, Nature Reviews-Drug Discovery, 17, 7840, 261–279, 2020, Springer-Nature

[3] M. Hoffmann et al., SARS-CoV-2 variants B.1.351 and B.1.1.248: Escape from therapeutic antibodies and antibodies induced by infection and vaccination, bioRxiv, doi: https://doi.org/10.1101/2021.02.11.430787, 10.110/2021.01.11.430787, 2021, Cold Spring Harbor Laboratory

[4] A. Rambaut et al., A dynamic nomenclature proposal for SARS-CoV-2 lineages to assist genomic epidemiology, Nature Microbiology, 5, 1403–1407 2020, Springer-Nature

[5] K. Murphy and C. Weaver, Janeway’s Immunobiology, 9th edition, 2017, Garland Science, Taylor & Francis Group, LLC

[6] R. Penner, Backbone Free Energy Estimator Applied to Viral Glycoproteins, Journal of Computational Biology, 27, 10, 1495–1508, 2020, Liebert

[7] R. Penner, Conserved High Free Energy Sites in Human Coronavirus Spike Glycoprotein Backbones, Journal of Computational Biology, 27, 11, 1622–1630, 2020, Liebert

[8] R. Penner et al., Hydrogen bond rotations as a uniform structural tool for analyzing protein architecture, Nature Communications, 5, 5803, 2014, Springer-Nature

[9] F.M. Pohl, Empirical protein energy maps, Nature New Biololgy 234, 277–279, (1971), Springer-Nature.

[10] A.V. Finkelstein, et al., Boltzmann-like statistics of protein architectures: Origins and consequences. In B.B. Biswas and S. Roy, eds., Proteins: Structure Function, and Engineering. Subcellular Biochemistry 24, 1–26, (1995), Springer, Boston, MA.

[11] A.V. Finkelstein, et al., Why do protein architectures have Boltzmann-like statistics? Proteins 23, 142–150, 1995, Springer-Nature.

[12] A.C. Walls et al., Structure, Function, and Antigenicity of the SARS-CoV-2 Spike Glycoprotein, Cell, 181, 2, 281–292, 2020, Cell Press

[13] R. Henderson et al., Controlling the SARS-CoV-2 spike glycoprotein conformation, Nature Structural and Molecular Biology, 27, 10, 925–933, 2020, Springer-Nature

[14] M. McCallum et al., Structure-guided covalent stabilization of coronavirus spike glycoprotein trimers in the closed conformation, Nature Structural and Molecular Biology, 27, 10, 942–949, 2020, Springer-Nature

[15] T. Zhou et al., Cryo-EM Structures of SARS-CoV-2 Spike without and with ACE2 Reveal a pH-Dependent Switch to Mediate Endosomal Positioning of Receptor-Binding Domains, Cell Host Microbe, 28, 6, 867–979, 2020, Cell Press

[16] C. Toelzer et al., Free fatty acid binding pocket in the locked structure of SARS-CoV-2 spike protein, Science, 370, 6517, 725–730, 2020, American Academy of Arts and Sciences

[17] A.G. Wrobel et al., SARS-CoV-2 and bat RaTG13 spike glycoprotein structures inform on virus evolution and furin-cleavage effects, Nature Structural and Molecular Biology, 27, 8, 763–767, 2020, Springer-Nature

[18] J. Juraszek et al., Stabilizing the closed SARS-CoV-2 spike trimer, Nature Communications, 12, 1, 244, 2021, Springer-Nature

[19] C. Xu et al., Conformational dynamics of SARS-CoV-2 trimeric spike glycoprotein in complex with receptor ACE2 revealed by cryo-EM, Science Advances, 7, 1, 5575, 2021, American Academy of Arts and Sciences

[20] S.M. Gobeil et al., D614G Mutation Alters SARS-CoV-2 Spike Conformation and Enhances Protease Cleavage at the S1/S2 Junction, Cell Reports, 34, 2, 108630, 2021, Cell Press

[21] R. Ganji and P.R. Hemachandra, Impact of COVID-19 on Mitochondrial-Based Immunity in Aging and Age-Related Diseases, Frontiers in Aging Neuroscience, 12, 502, 2021, Frontiers Research Foundation.

[22] C.O. Barnes, et al., SARS-CoV-2 neutralizing antibody structures inform therapeutic strategies, Nature, 588, 7839, 682–687, 2020, Springer-Nature.

[23] Z. Liu et al., Identification of SARS-CoV-2 spike mutations that attenuate monoclonal and serum antibody neutralization, Cell Host and Microbe, 29, 477–488, 2021, Elsevier Inc..

